# Effect of levodopa on human brain connectome in Parkinson’s disease

**DOI:** 10.1101/2021.06.11.448038

**Authors:** Sajjad Farashi, Mojtaba Khazaei

**Affiliations:** Autism Spectrum Disorders Research Center, Hamadan University of Medical Sciences, Hamadan, Iran; Dental Implant Research Center, Hamadan University of Medical Sciences, Hamadan, Iran; Department of Neurology, School of Medicine, Sina (Farshchian) Educational and Medical Center, Hamadan University of Medical Sciences, Hamadan, Iran

**Keywords:** Parkinson’s disease, Levodopa, L-dopa, EEG, Graph theory

## Abstract

**Objective:** Levodopa-based drugs are widely used for mitigating the complications induced by PD. Despite the positive effects, several issues regarding the way that levodopa changes brain activities have remained unclear.

**Methods:** A combined strategy using EEG data and graph theory was used for investigating how levodopa changed connectome and processing hubs of the brain during resting-state. Obtained results were subjected to ANOVA test and multiple-comparison post-hoc correction procedure.

**Results:** Results showed that graph topology of PD patients was not significantly different with the healthy group during eyes-closed condition while in eyes-open condition statistical significant differences were found. The main effect of levodopa medication was observed for gamma-band activity of the brain in which levodopa changed the brain connectome toward a star-like topology. Considering the beta subband of EEG data, graph leaf number increased following levodopa medication in PD patients. Enhanced brain connectivity in gamma band and reduced beta band connections in basal ganglia were also observed after levodopa medication. Furthermore, source localization using dipole fitting showed that levodopa prescription suppressed the activity of collateral trigone.

**Conclusion:** Our combined EEG and graph analysis showed that levodopa medication changed the brain connectome, especially in the high-frequency range of EEG (beta and gamma).

**Highlights:**

- No differences were found between graph features between ON and OFF PD cases in eyes-closed.
- Levodopa enhanced connectivity in gamma band in PD patients.
- Levodopa inhibited brain connectivity in beta band.

## 1. Introduction

Parkinson’s disease (PD) is one of the main neurodegenerative diseases that mostly observed in elderly people. The deterioration of dopaminergic neurons in basal ganglia especially in the substantia nigra pars compacta region degrades the dopamine level in the brain. This causes several drawbacks including movement disorder (resting tremor, freezing of gait, postural tremor, rigidity, bradykinesia and disturbed gait dynamics ^1^) and also many other non-motor disorders like cognitive impairment, sleep disorder, smell disorder and so on ^2^. Although there is no exact treatment for PD, the levodopa-based medication (L-dopa medication) is the well-accepted choice for mitigating complications due to PD. L-dopa based drugs can pass through the blood-brain barrier, be synthesized to dopamine and in this way compensate for the dopamine insufficiency to some extent ^3^. Even though L-dopa therapy shows positive effects for some PD patients, it is not clear how this drug affects brain activity ^3^.

The effect of L-dopa medication on the electrical activity of the brain in PD patients was the subject of several studies. Jackson et al. revealed that during resting-state, beta waves had sharper profiles in the absence of drug administration ^4^. They also showed that EEG amplitude-phase coupling in high-frequency content of EEG (i.e. beta and gamma bands) was increased when the medication was terminated. Furthermore, without anti-Parkinsonism drugs, brain activities showed asymmetric patterns while levodopa prescription caused more symmetric activity ^4^. The reduced amount of beta wave synchrony in the motor cortex following drug prescription was another consequence that was suggested as the reason for improved motor functions. Lofredi et al. (2018) reported that following depletion of dopamine in brain regions, a significant decrease of gamma wave power and gamma bursting pattern were observed. The vanished gamma wave contribution in EEG power might be the reason for bradykinesia in PD patients ^5^. Miller et al. used EEG extracted biomarkers for PD patients in L-dopa prescription state (ON-medication) and termination of L-dopa medication state (OFF-medication) to check how drug prescription changed the relationship between EEG and movement impairments ^6^. The results showed that both coherence and amplitude-phase coupling were higher for OFF-medication state. Decreased coherence and amplitude-phase coupling were correlated with improvement in movement capability in PD patients ^6^. Stegemoller et al. investigated EEG power spectrum in alpha and beta bands for ON and OFF medication states when PD subjects moved their fingers. They showed that damping of alpha and beta power was observed for healthy, ON-medication and OFF-medication PD patients just before finger movements, however, the damping ratio for faster finger movements was higher for OFF-medicated patients ^7^. Using EEG time-series George et al. showed that during resting-state the pair-wise coherence in the beta band was higher for PD patients in OFF-medication state. An over-synchronization was also observed for PD patients compared with the healthy group that was reduced following levodopa prescription ^8^.

Graph theoretical approaches have gained special attention for analyzing brain activities and especially its connectome in PD. Reduced functional connection in sensory-motor network and supplementary motor area in OFF-medication state as compared with healthy subjects was reported by Schneider et al. ^9^. It was shown that L-dopa medication enhanced functional connectivity in the above-mentioned areas. Utianski et al. used graph networks constructed from EEG time-series of PD patients with dementia, PD without dementia and healthy subjects to investigate if brain connectivity was different between these groups. This study showed that local integration in all frequency bands was better for healthy and PD patients without dementia, especially in the alpha frequency band ^10^. Graph analysis also showed a decrease in local clustering following disease progression ^11^. Reduced functional connectivity, when connectivity was calculated using phase lag index, was reported to be correlated with disease severity ^12^. It was proposed by some other studies that reduced brain connectivity in alpha band might be related to cognitive impairments in PD patients ^12,13^.

In this study, the main focus was on the way that levodopa medication influences brain function during resting-state. Resting-state is a condition in which the activity of the brain is evaluated in the absence of any stimulus. Usually, resting-state can be divided into three distinct conditions including eyes-closed (EC), eyes-open (EO) and eyes-open with fixed gaze ^14^. Studies revealed significant differences in brain dynamics during EO and EC conditions ^15,16^. According to significant differences between brain activities during EO and EC conditions, it is important to check the effect of L-dopa medication on EO and EC resting-state conditions separately. Here, it was hypothesized that L-dopa medication changed not only brain connections, also the activation of brain areas should be altered.

In the current study, two different resting-state conditions (i.e. EO and EC) were considered to investigate how L-dopa medication influences brain connectivity in PD patients. Furthermore, based on the graph network analysis, the main brain areas engaged in L-dopa medication were discussed. For these purposes, EEG time-series during resting-state and a graph theoretical approach were used. A minimum spanning tree (MST) topology was constructed for the EEG data of each person and further analyses were performed according to this graph.

## 2. Material and Methods

### 2.1 Dataset description

In this study, a publicly available EEG dataset from the University of New Mexico was used ^17^(available at: http://predict.cs.unm.edu/downloads.php, Project Name: Parkinson’s Rests). This dataset contains the brain waves for 27 PD patients in two distinct conditions (ON-medication when PD patients are prescribed dopaminergic drugs and OFF-medication when the same PD patients withdraw anti-parkinsonism drugs for 15 hours). Furthermore, EEG recordings for the age and sex-matched normal participants (n=27) are available in this dataset. PD participants visited the recording lab twice at a 7-day interval. Half of the patients were in ON medication at the first visit, while others were in OFF state. All EEG recordings were performed at 9:00 AM for all participants. Also, for PD group, several behavioral tests including Mini Mental State Exam (MMSE), Beck Depression Inventory (BDI) test, the North American Adult Reading Test (NAART) and Unified Parkinson’s Disease Rating Scale (UPDRS) motor score test were performed. All selected PD patients had MMSE scores higher than 26. EEG recordings (10-20 international system) were performed using a 64-channel system with the sampling frequency of 500Hz while simultaneously EMG activity produced by vertical eye movements and the acceleration of hand movements using an accelerometer system were recorded for each participant. Recording included 1-min eyes-closed (EC) at the resting-state condition and followed by 1-min eyes-open (EO) at resting-state condition while subjects were in a relaxed position. The start and end time points for EC and EO conditions were specified on recorded time-series as suitable labels.

### 2.2 Preprocessing step

The dataset was imported to EEGLAB toolbox ^18^ of MATLAB 2017 (Mathworks, MA, USA). In the first preprocessing step unnecessary channels like vertical eye movements and hand acceleration data were removed. The baseline value for each remained channel was removed using a moving average strategy, then for canceling data outside the EEG frequency band, FIR filtering in 0.5-50 Hz was performed. FIR filtering with linear phase guaranteed that time-series distortion was avoided. For removing artifacts and noises with the overlapped frequency content with EEG time-series such as EMGs, a blind source separation strategy using independent component analysis (ICA) with fastICA algorithm was performed and the component contained most of non-EEG contents were removed. The denoised data were investigated visually to be sure that all non-EEG artifacts were removed from the data. For denoised data, bad channels were found using kurtosis measure and a Z-score of 5 and were interpolated by neighboring channels in a 10-20 system. Here, no re-referencing was done and the original reference and ground of the dataset (CPz and AFz, respectively) were used. The preprocessed data was segmented to EEG for eyes-closed and EEG for eyes-open segments for further analyses. Analyses in EEG subbands were conducted by filtering pre-processed EEG time-series in 0.5-3.5 Hz (Delta band), 4-8 Hz (Theta band), 8-12 Hz (Alpha band), 12-30 Hz (Beta band) and 30-50 Hz (Gamma band).

### 2.3 MST graph and features

The pre-processed data for each subject was used for constructing a minimum spanning tree (MST) graph for that subject. MST is an acyclic graph that its edges pass all nodes of the system while the minimum total edge weight is satisfied. Here nodes are recording electrodes (or equivalently brain surface regions beneath each electrode) and edges are the possible connections between nodes that might be due to the anatomical or functional connection. MST eliminates the bias problem during the comparison of networks which exists for traditional graph methodologies ^19^.

In order to investigate how different brain regions are connected, the association between EEG time-series acquired by electrodes placed on the scalp can be calculated using different methods such as correlation or weighted phase lag index (WPLI) which the latter is robust against the volume conductance property of the brain ^20^. From the constructed MST graph using WPLI several features including graph diameter, radius, betweenness centrality (BC), mean eccentricity (mEC), hierarchical tree (HT), degree and leaf number were extracted. The description for each measure was given in Table 1.

**Table 1.** MST graph extracted features.

| **Measure** | **Description** |
| --- | --- |
| Leaf number | Number of nodes connected to only one another node and is defined for the whole network. |
| Node degree | It is defined for each node and determines how many connections exist between that node and other nodes of the network. |
| Diameter | It is the maximum possible distance between two vertices in the graph <sup>21</sup>. |
| Radius | It is the minimum of maximum distances between a node and other nodes. |
| Betweenness centrality(BC) | It is defined for each node and determines how many times the node is placed on the shortest path between all other pairs of nodes. |
| Mean Eccentricity (mEC) | It is defined for each node as the maximum distance between that node and all other nodes of the system. A mean value for all nodes defines the mean eccentricity of the graph. |
| Hierarchical Tree (HT) | It shows the hierarchical structure of a MST. |

Higher BC value indicates the hubness. Hubs are the main information processing units in the network and act as communication ports in the system. For each MST network, the maximum value was considered as BC measure of the graph. HT is dependent on the graph structure and determines the sub-trees children of a parent node. Lower HT values indicate line-like trees while for star-like topologies higher HT is obtained. In a hierarchical structure, hubs are connected to nodes that have lower degrees ^22^.

### 2.4 Statistical analysis

The distribution of graph measures was investigated using Shapiro-Wilk goodness-of-fit test ^23^ to be sure that data come from a normal distribution. The significance of the difference between graph extracted features of PD group (in ON and OFF-medication states) and healthy individuals were tested according to ANOVA. For finding between-group differences and corrected the p-values, post-hoc analysis using multiple-comparison correction strategy (Fisher’s least significant difference procedure) was performed ^24^. All analyses were performed using the statistical toolbox of MATLAB 2017 (Mathworks, MA, USA). Furthermore, the correlation between MST graph measures in different frequency bands and UPDRS motor scores were obtained using Pearson’s correlation and the significance of the obtained correlation was assessed using the student t-test. The significant level for accepting statistical significant difference was 0.05

## 3. Results

In this study, a comparison between PD subjects in ON and OFF medication states and matched healthy individuals was performed using extracted features of MST graph on basis of an EEG dataset. In Table 2, statistically significant differences between these groups in EEG subbands were reported. The only significantly different feature between ON and OFF-medication conditions was graph leaf number in the beta frequency band, where L-dopa medication decreased graph leaf number in PD patients.

**Table 2.** Statistically significant differences between MST features for ON-medication (ON), OFF-medication (OFF) and healthy (H) groups in different frequency contents of EEG waves. CIL and CIH are lower and upper mean difference values for 95% confidence intervals, respectively. P-values are obtained according to the multiple comparison correction test.

| Groups | Feature | Resting condition | Frequency | Mean difference | CIL | CIH | P value |
| --- | --- | --- | --- | --- | --- | --- | --- |
| ON,H | Eccentricity | EO | Theta | -0.57 | -1.13 | -0.01 | 0.046 |
| ON,H | Diameter | EO | Alpha | -1.94 | -3.11 | -0.78 | 0.001 |
| OFF,H | Diameter | EO | Alpha | -1.44 | -2.60 | -0.27 | 0.016 |
| ON,H | Eccentricity | EO | Alpha | -1.34 | -2.14 | -0.53 | 0.001 |
| OFF,H | Eccentricity | EO | Alpha | -1.00 | -1.81 | -0.19 | 0.015 |
| ON,H | Radius | EO | Alpha | -0.98 | -1.57 | -0.39 | 0.001 |
| OFF,H | Radius | EO | Alpha | -0.73 | -1.31 | -0.14 | 0.015 |
| **ON,OFF** | **Leaf number** | **EO** | **Beta** | **5.26** | **1.79** | **8.72** | **0.003** |
| ON,H | Diameter | EO | Gamma | -0.04 | -0.09 | -0.003 | 0.036 |
| ON,H | Eccentricity | EO | Gamma | -0.04 | -0.07 | -0.002 | 0.037 |
| ON,H | Radius | EO | Gamma | -0.02 | -0.05 | -0.001 | 0.035 |

Previous studies according to fMRI data revealed that brain regions at rest were functionally integrated over multiple frequency bands ^25^. In this perspective, it seems that for a better conclusion about the role of different brain regions in a specific task, wider or multiple frequency bands should be considered. In Table 3 differences between MST graph features between PD (ON and OFF-medication conditions) and healthy individuals for lower frequency content of EEG (delta+ theta) and upper frequency content of EEG (alpha+ beta and beta+gamma) as well as the whole frequency range of EEG were reported.

**Table 3.** Statistically significant differences between MST features of ON-medication (ON), OFF-medication (OFF) and healthy (H) groups for different frequency contents of EEG waves. CIL and CIH are lower and upper value for 95% confidence intervals, respectively. P-values are obtained according to post-hoc multiple comparison correction test.

| Groups | Feature | Resting condition | Frequency | Mean difference | CIL | CIH | P value |
| --- | --- | --- | --- | --- | --- | --- | --- |
| **Low-frequency content** |  |  |  |  |  |  |  |
| ON,H | Diameter | EO | Delta+Theta | -0.72 | -1.13 | -0.32 | <0.001 |
| OFF,H | Diameter | EO | Delta+Theta | -0.46 | -0.87 | -0.06 | 0.027 |
| ON,H | Eccentricity | EO | Delta+Theta | -0.57 | -0.89 | -0.25 | <0.001 |
| OFF,H | Eccentricity | EO | Delta+Theta | -0.36 | -0.68 | -0.04 | 0.027 |
| ON,H | Radius | EO | Delta+Theta | -0.38 | -0.59 | -0.17 | <0.001 |
| OFF,H | Radius | EO | Delta+Theta | -0.24 | -0.45 | -0.03 | 0.023 |
| **High-frequency content** |  |  |  |  |  |  |  |
| **ON,OFF** | **Leaf Number** | **EO** | **Alpha+Beta** | **3.67** | **0.04** | **7.29** | **0.047** |
| OFF,H | Leaf Number | EO | Alpha+Beta | -3.85 | -7.48 | -0.23 | 0.037 |
| ON,H | Hierarchy | EO | Beta+Gamma | -0.03 | -0.06 | -0.00 | 0.044 |
| OFF,H | Hierarchy | EO | Beta+Gamma | -0.03 | -0.06 | -0.00 | 0.017 |
| **Whole frequency range** |  |  |  |  |  |  |  |
| **ON,OFF** | **Diameter** | **EO** | **0.5-50 Hz** | **-0.04** | **-0.07** | **-0.00** | **0.035** |
| OFF,H | Diameter | EO | 0.5-50 Hz | 0.04 | 0.00 | 0.07 | 0.036 |
| **ON,OFF** | **Eccentricity** | **EO** | **0.5-50 Hz** | **-0.03** | **-0.06** | **-0.00** | **0.027** |
| OFF,H | Eccentricity | EO | 0.5-50 Hz | 0.03 | 0.00 | 0.06 | 0.030 |
| **ON,OFF** | **Radius** | **EO** | **0.5-50 Hz** | **-0.02** | **-0.04** | **-0.00** | **0.029** |
| OFF,H | Radius | EO | 0.5-50 Hz | 0.02 | 0.00 | 0.04 | 0.027 |

According to Table 3, graph diameter, radius and eccentricity were significantly different between PD patients and healthy individuals when lower frequency content of EEG was considered. Results of Table 3 for lower frequency content of EEG indicated smaller graph diameter, radius and eccentricity for PD patients compared with healthy individuals, however, ON-medication increased such differences. In the higher frequency region of EEG (alpha + beta), leaf number of MST graph was significantly higher for ON-medication state compared with OFF-medication. Furthermore, in beta+gamma band the hierarchical property of PD patients was lower for healthy individuals, even though the confidence intervals showed the differences were negligible.

Statistical analysis (Tables 2 and 3) revealed that the graph topology was different for EO condition between PD (ON/OFF-medication) and healthy groups and no statistically significant differences were found between groups in EC condition. In this regard, all reported results of this study were limited to EO condition. The brain connectivity patterns when different frequency contents of EEG were considered were shown in Figs 1 and 2. In these figures, the strongest connection weight between pairs of electrodes in three conditions (i.e. ON, OFF and healthy conditions) was obtained and connections with weight lower than one-third of that strongest weight were preserved and looser connections were removed. This was performed to find the most important connected areas.

**Fig 1.**
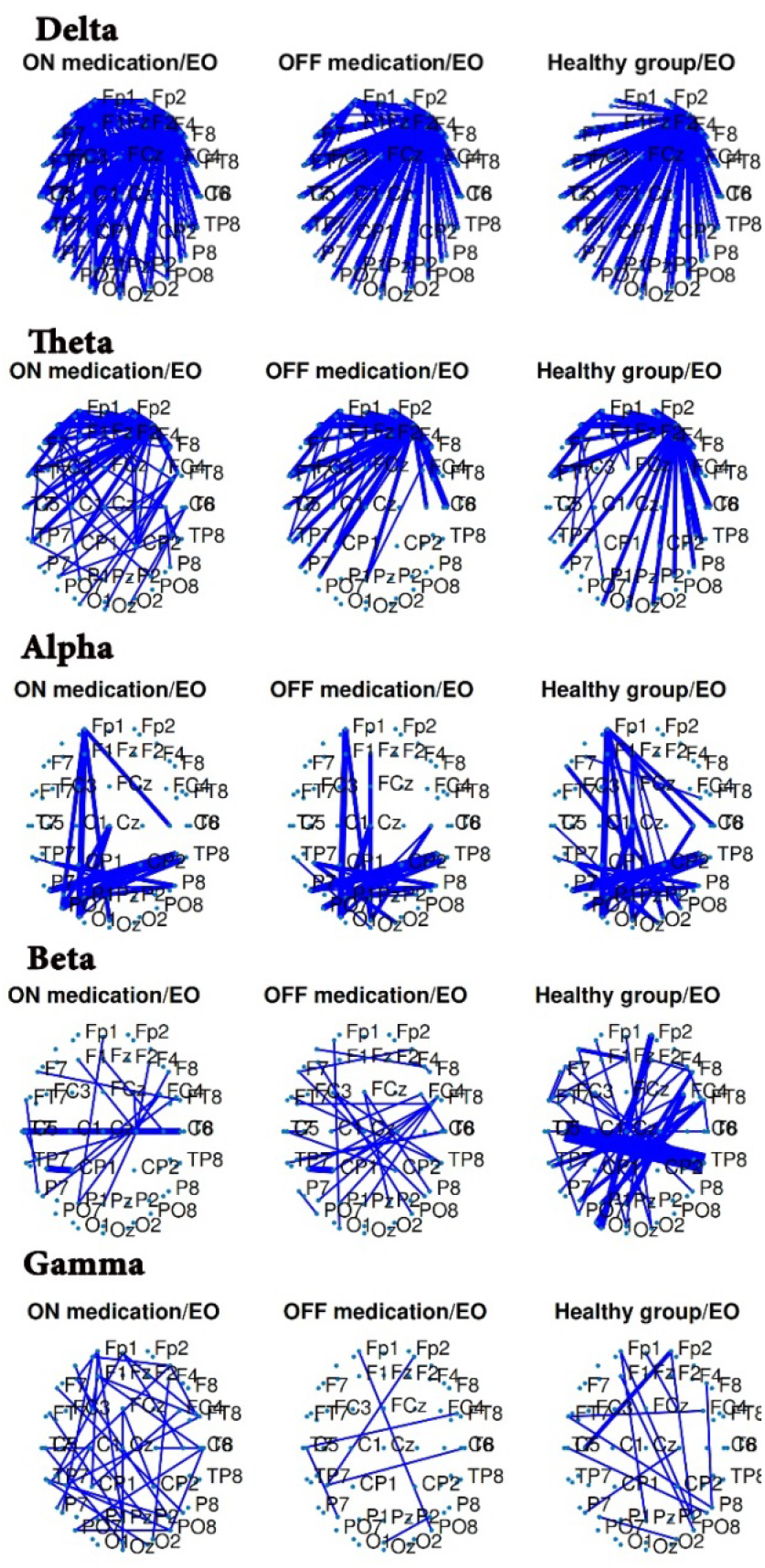
Connections between different brain areas in EO resting-state for ON-medication PD (first column), OFF-medication PD (second column) and healthy (third column) subjects in EEG specific bands (delta, theta, alpha, beta and gamma). Only strong connections were preserved.

**Fig 2.**
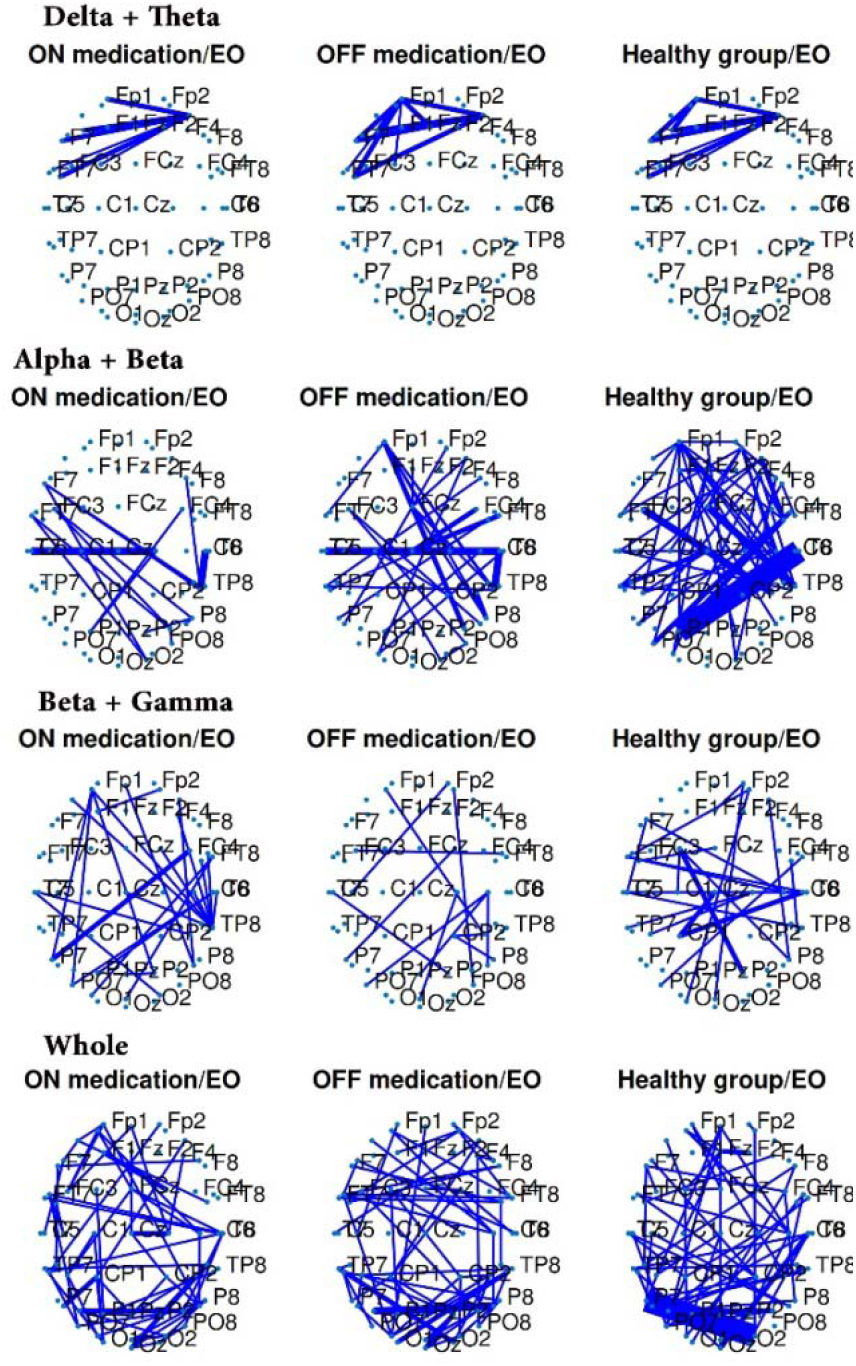
Connections between different brain areas in EO resting-state for ON-medication PD (first column), OFF-medication PD (second column) and healthy (third column) subjects when low-frequency (Delta+Theta), high-frequency (Alpha+Beta and Beta+Gamma) and whole frequency content of EEG was considered. Only strong connections were preserved.

BC is a measure of hubness for brain processing units. It shows that how different brain areas contribute to information processing during a specific task. Since the main objective of this study was the effect of L-dopa medication on brain activity in resting-state, the difference between BC measure in ON-medication and OFF-medication conditions during resting-state were shown. According to Fig 3, statistical analyses showed that brain hubness was different in temporoparietal (beta and beta+gamma and whole frequency bands) and parietal (gamma band) between ON and OFF-medication conditions. A dipole fitting strategy (see Fig 4) was used to investigate what kind of sources produced such differences.

**Fig 3.**
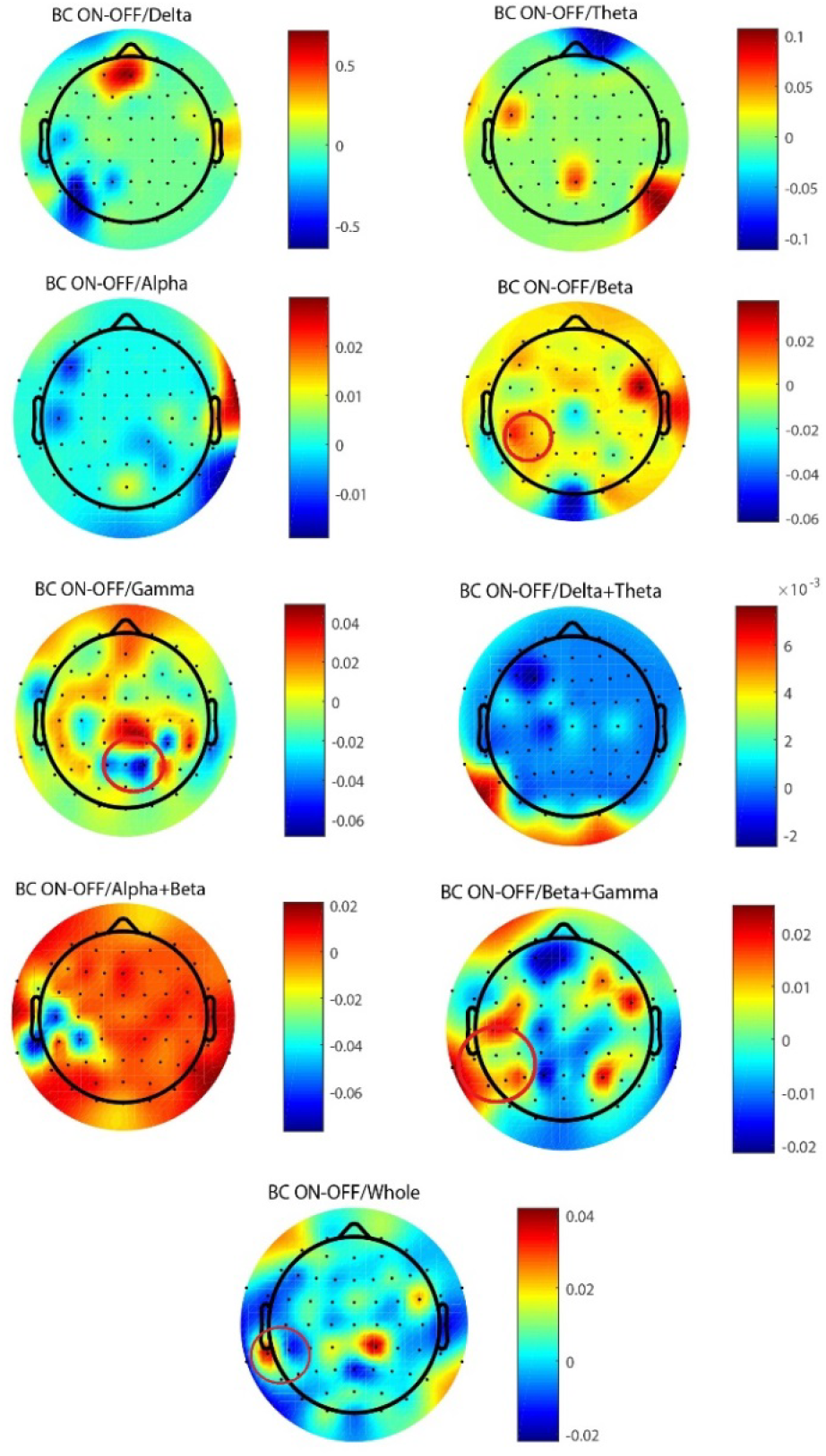
Topoplots for difference of BC measure between ON and OFF medication states during EO resting-state. Each figure had its own colorbar scale. Red circles showed brain locations where statistical significant different BC measures between ON and OFF states were found.

**Fig 4.**
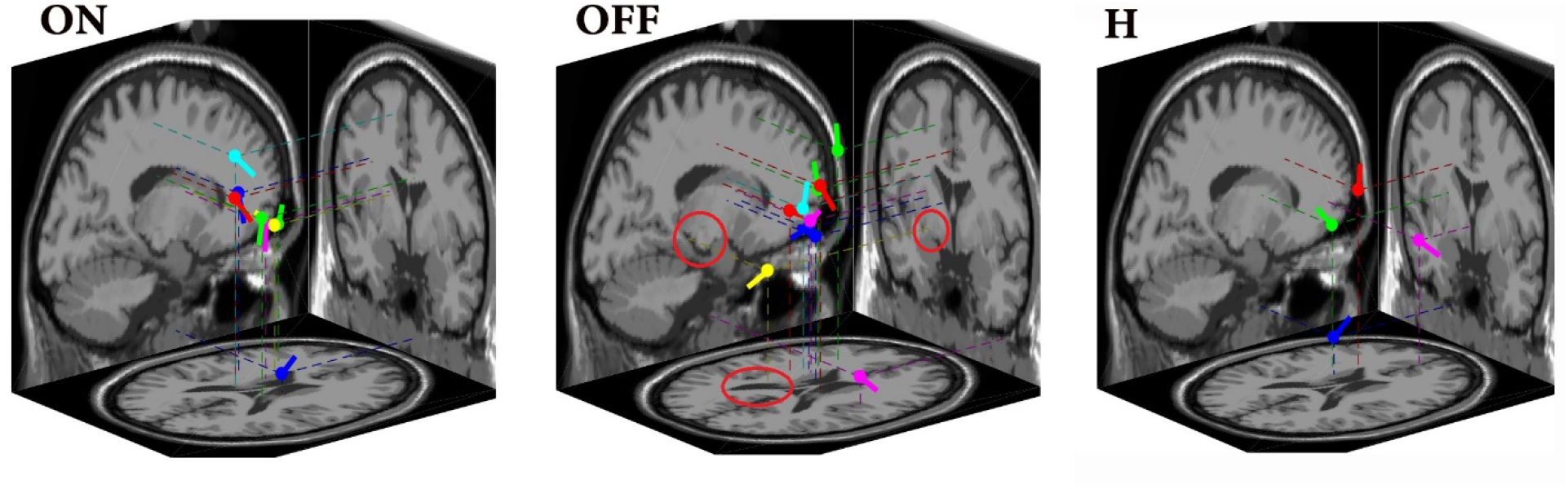
Dipole fitting for EO resting-state condition for ON-medication (ON), OFF-medication (OFF) and healthy (H) groups. Dipoles were the most frequent between cases in each group.

According to statistical analysis, some MST graph features were significantly different between ON and OFF medication for PD patients. These features were graph leaf number (in beta and alpha+ beta bands), diameter, radius and eccentricity when total frequency range of EEG time-series was considered. It was important to check if such brain connection alteration due to L-dopa medication had positive effects on motor activity in PD patients. For this purpose, the correlation between alteration of MST graph characteristics and the alteration of UPDRS motor section scores were calculated and the results were depicted in Table 4.

**Table 4.** Correlation analysis for MST graph extracted features change and UPDRS motor section following L-dopa medication.

| Feature | Correlation | P value | Frequency |
| --- | --- | --- | --- |
| Diameter | 0.493 | 0.008 | whole |
| Radius | 0.497 | 0.008 | whole |
| Eccentricity | 0.495 | 0.008 | whole |
| Leaf number | -0.429 | 0.026 | Beta |
| Leaf number | -0.503 | 0.007 | Alpha+ Beta |

## 4. Discussion

For EO resting-state condition in theta, alpha and gamma bands, after L-dopa medication, PD patients had lower graph eccentricity value as compared with the healthy group. The diameter and radius of MST graph of PD patients after drug administration in alpha and gamma were also lower in ON-medication PD patients as compared with the healthy group. From Table 2, it was obvious that the graph eccentricity, radius and diameter in the alpha band with or without anti-parkinsonism drug administration (OFF-medication) were lower for PD patients compared with healthy individuals. This result showed that PD was the responsible factor for graph characteristic change in the alpha band. However, when the mean differences and confidence intervals were considered, it was obvious that L-dopa medication-induced larger differences between healthy and PD participants in the alpha band.

Graph diameter is a measure for system integration capacity ^26^ and determines the efficiency of information exchange in the whole system. Low diameter graph is related to a more efficient processing system when remote regions are collaborating for performing a task ^27^. In higher diameter graphs, it is expected that a message takes more steps to reach the other end of the network, especially if the leaf number be lower ^28^. In the case of low diameter and low eccentricity values, the system might rely highly on special hub nodes (i.e. a star-like network ^29^). In this regard, according to Tables 2 and 3, in alpha, gamma and delta+ theta bands, since L-dopa medication reduced both diameter and eccentricity, it could be concluded that L-dopa medication increased the hubness of the brain system. While in alpha and delta+theta bands the increased hubness seemed to be intrinsic to PD (i.e. for both ON and OFF-medication cases higher hubness was observed compared with healthy individuals), L-dopa medication especially changed brain connection topology toward a star-like shape in the gamma band.

In overall, the results (Table 2) showed that when the alpha rhythm of EEG was considered, the brain topology of PD patients was more similar to a star-like network during EO resting-state condition, as compared with the healthy group. The L-dopa medication reinforced such star-like property in alpha band, however, no statistically significant difference was found between ON and OFF-medication states. Furthermore, in the gamma band, levodopa medication changed the MST graph topology toward a star-like shape. When multiple-frequency bands were considered, it was observed that in the lower frequency content of EEG (delta+ theta), like the results obtained for alpha band (Table 3), MST graph topology related to PD cases was closer to star-like shape (i.e. lower graph diameter, radius and eccentricity), while this closeness was more significant after L-dopa medication. This result showed that in lower-frequency range of EEG, as well as alpha frequency range, the brain global efficiency was higher for PD cases as compared with healthy individuals. This result was consistent with the fact that EEG activities were slowed during PD progress ^30^. In higher frequency ranges of EEG (beta+ gamma), the brain for PD cases had lower hierarchical processing property (p<0.05) that it was not influenced by L-dopa medication. A hierarchical metric is used to evaluate the trade-off between large-scale integration and the overload of the central node in the MST.

According to Table 3, when lower frequency content of EEG was considered, differences between PD and healthy groups existed for both ON and OFF-medication conditions and no significant differences were found between ON and OFF states. This outcome indicated that differences were not due to medication, instead such differences might be due to disease.

When brain connections in distinct frequency bands were investigated (Table 2), it was clear that MST graph topological features remained almost unchanged between ON and OFF medication conditions. The only difference between ON and OFF medication states was observed in beta band for leaf number, where the leaf number was significantly higher for ON medication state. Larger leaf numbers in ON medication compared with OFF medication state in the beta frequency band might indicate functional connections in the brain altered in a way that more similar star-like topology manifested in the network. This possibly indicated the increase in global efficiency upon L-dopa medication for PD patients. When multiple frequency bands were considered, in low-frequency region (Delta+ Theta), no significant difference between ON and OFF-medication was observed, while for higher frequency content of brain electrical activities (alpha+ beta), graph constructed according to brain connections for ON-medicated patients showed higher leaf numbers that could be interpreted as a topology more similar to star-like topologies. Comparing Tables 2 and 3, it seemed that leaf number difference in alpha + beta band was mainly due to brain activities in beta instead of alpha band. Leaf number increase following L-dopa medication was significantly correlated with a decrease in UPDRS motor section (see Table 4). This indicated that increasing processing units and changing the pattern of the brain connectome toward a star-like topology might result in better movement actions following L-dopa medication. Considering whole frequency range of EEG (0.5-50 Hz), statistically different graph measures between ON and OFF medication states were eccentricity, diameter and radius, however, the difference seemed to be very small (95% CI for mean difference:[−0.06, 0], [−0.07, 0] and [−0.04, 0], respectively). By comparing Tables 2 and 3 for subband analyses and analysis in the whole-frequency range of EEG, it could be concluded that PD had multi-frequency effects on brain activities.

From Fig 1, when low-frequency content of EEG was considered (delta and theta bands), the main driver of the brain network was supposed to be in the frontal cortex during EO resting-state for both PD and healthy cases. Frontal lobe is an important unit for visual perception and visually guided behavior ^31^ during EO condition. Theta frequency oscillations in the frontal area and anterior cingulate cortex are expected during eyes-open due to the brain attentional processes ^32^. According to Fig 1, L-dopa medication created a strong hub in the frontal area compared with OFF-medication and healthy conditions (see Fig 1, Delta ON-medication EO and Fig 3, BC ON-OFF/Delta). However, statistical analysis revealed that such an effect was not significant (p>0.05) between BC measures of ON and OFF-medication groups. In the alpha band, a clear connection loss between frontal and posterior regions was observed as compared with healthy individuals (Fig 1, third row). In humans, basal ganglia modulates the connectivity between the prefrontal cortex and posterior region during shift attention in EO state. Since in PD the most infected region is basal ganglia, the lost association between frontal and posterior regions in PD cases in Fig 1 in the alpha band might be due to the basal ganglia dysfunction in PD patients ^33^. Fig 3 (BC ON-OFF/alpha) showed that two distinct brain locations in fronto-temporal area were active during the OFF-medication condition compared with the ON-medication, even though statistical analysis did not confirm their significance.

During resting-state, overexpression of STN beta activity that is attenuated with starting of voluntary movement is observed ^34^. Previous studies showed the positive effect of levodopa medication on normalizing brain connectivity by increasing beta coherence ^35^. It was consistent with obtained results of this study in which L-dopa medication reduced the strong connections during EO resting-state (Fig1, fourth row, ON medication/EO as compared with OFF medication EO).

When multi-frequency spectrum of EEG was considered (Fig 2), the enhanced connection in the high-frequency range of EEG (beta + gamma) and reduced over-expression of central region (possibly related to basal ganglia) in alpha+beta band were observed.

According to Fig 3 in the beta band, following L-dopa medication BC measure in the left temporo-parietal region was significantly different between ON and OFF states (Fig 3, BC ON-OFF/Beta panel, circles). The nested network of brain connectivity for gamma band in OFF-medicated PD patients as compared with healthy individuals was observed in Fig 1(see Fig 1, Gamma OFF medication/EO panel). Increased gamma power is associated with increased motor activity and arousal levels, which are both degraded in PD patients ^36^. Also, dopamine depletion in the brain reduced gamma wave contribution in brain functioning ^5^. In this regard, reduced gamma band activity and connectivity might be the reason for such a nested network. Furthermore, in animal models of PD, it was shown that following the L-dopa medication dyskinesia could be induced that was associated with increased gamma oscillatory power ^37^. The increased gamma band connectivity was observed in Fig 1 (Gamma ON medication/EO). As Fig 3 (BC ON-OFF/Gamma) showed BC difference between ON and OFF states appeared at the posterior region of the head model. Statistical analysis showed that a significant difference (p=0.045) existed at PO4 electrode location in which a reduced BC measure was observed following L-dopa medication. According to Fig 4, when dipole fitting of ON and OFF-medication states were compared, it was clear that more brain sources in mid-brain area were activated during OFF-medication state as compared with ON-medication state. For both ON and OFF-medication PD cases, this extra activity in the mid-brain area (i.e. over-activation of basal ganglia ^38^) was obvious in comparison with healthy subjects. L-dopa medication reduced over-activation of the basal ganglia region, since fewer numbers of dipoles were produced in the mid-brain following medication. Furthermore, an extra source of activity in collateral trigone region appeared in OFF-medication state (circles in Fig 4, OFF panel) that might produce the observed hub in the gamma band (Fig 3, BC ON-OFF/Gamma). The collateral trigone is a triangular area at the intersection between temporal and occipital branches of the lateral ventricle. The projection of dipoles in the left hemisphere for OFF-medication case might be the reason for different hubness in the temporo-parietal region observed for beta, beta+gamma and whole frequency bands (see Fig 3, BC ON-OFF/Beta, Beta+Gamma and Whole panels).

## 5. Study limitation

In this study, the sample size for each group (healthy, ON-medication and OFF-medication) was 27. Even though this sample size was larger than many other similar researches, a bigger sample size could obtain statistical analysis with a higher power. Furthermore, the time duration for EEG time-series for EO and EC conditions was 1-minute. A longer duration might discover long-time interactions between brain regions following L-dopa medication.

## 6. Concluding remarks

The results of this study showed that the brain connectome might be affected by PD. In addition, L-dopa medication also affected the topological properties of brain connections. Increasing leaf number following L-dopa medication was observed when beta or alpha+beta content of EEG were considered. This increased leaf number was significantly associated with enhanced movement ability. The sparse Brain connection in the gamma frequency band was observed for PD cases that were compensated by L-dopa medication. Furthermore, L-dopa medication regulated brain connections in the beta frequency band toward suppression of over-activation activities of brain, especially in the basal ganglia region. Perhaps in this way L-dopa medication mitigates some type of movement disorders in PD patients. In addition, L-dopa medication highlighted some processing units (hubs) in temporal and temporo-parietal regions observed at beta+ gamma and gamma frequency band of EEG, while suppressed a processing unit in the posterior region observed at beta band of EEG. Dipole fitting showed that the activity of the collateral trigone region, a deep area in the brain, might be responsible for such different hubness property during ON and OFF-medication conditions. From a topological perspective, this study showed that L-dopa medication changed brain topology toward a star-like structure especially when the high-frequency content of EEG was considered.

## Declarations

### Acknowledgments

Author would like to thank Deputy of Research and Technology, Hamadan University of Medical Sciences for its support for the current work.

### Ethics approval and consent to participate

Not applicable

### Consent for participate

Not applicable.

### Availability of data and materials

Not Applicable.

### Consent for publication

Not applicable

### Code availability

Not Applicable.

### Authors’ contributions

S.F: design, data analysis, wrote the manuscript; M.K: data analysis and wrote the manuscript.

